# Disruption of the surfactant protein A receptor SP-R210_L_ (CD245α/MYO18Aα) alters respiratory function and iron sequestration in alveolar macrophages of aged mice

**DOI:** 10.1101/2021.04.14.439860

**Authors:** Eric Yau, Todd M Umstead, Raz Abdulqadir, Kristin Fino, Zhiwei Guan, Sanmei Hu, Susan DiAngelo, Kavitha Hassan, Sarah S. Bingaman, Hannah Atkins, Timothy K Cooper, Amy C. Arnold, E. Scott Halstead, Zissis C. Chroneos

## Abstract

Previous studies demonstrated that the host defense collectins, surfactant protein A and complement component 1q, modulate tissue-dependent macrophage activation, pathogen clearance, and regulatory macrophage functions through the receptor SP-R210, which consists of two isoforms SP-R210_L_ and SP-R210_S_. These isoforms are encoded by alternatively spliced mRNAs of the *Myo18a MYO18A* gene in mice and humans. The present study in conditional transgenic mice revealed novel age-related functions of the SP-R210_L_ isoform in modulating pulmonary mechanics, iron sequestration in alveolar macrophages (AMs), and life-long maintenance of the alveolar macrophage population. Our findings support the novel idea that SP-R210_L_-deficient AMs undergo bi-directional epigenetic adaptation that results in chronic dysregulation of broncho-alveolar function, immune homeostasis, and maintenance of oncotic balance at the airway-capillary interface. Disruption of SP-R210_L_ increases the risk for development of severe interstitial lung disease during development and aging.

## INTRODUCTION

The macrophage-expressed isoforms of myosin XVIIIA (MYO8Aα and MYO18Aβ), named as SP-R210_L_ and SP-R210_S_, respectively, mediate immune and host defense functions as shared receptors for surfactant protein A (SP-A) and complement component 1q (C1q) in alveolar and peritoneal macrophages, respectively, as well as autonomous modulation of macrophage phenotype (1–6). SP-R210 is identical to the previously orphan antigen CD245 (7). Alternative splicing of *Myo18A* generates multiple protein isoforms categorized broadly as MYO18Aα, MYO18Aβ, and MYO18Aγ. MYO18Aα and MYO18Aγ are distinguished from MYO18Aβ by different amino-terminal PDZ domain and polyproline extensions, respectively and an additional carboxy-terminal tail in MYO18Aγ(8). Myo18Aα is more widely expressed, although not in the liver, heart, or skeletal muscle (9). MYO18Aβ is found in hematopoietic tissues, whereas MYO18Aγ is exclusive to heart and skeletal muscle (8–10). SP-R210_L_ occurs in mature macrophages, whereas SP-R210_S_ occurs in both mature and immature myeloid cells (9, 10). SP-A expression outside the lung and during pregnancy mediates immunological surveillance and physiological control of uterine smooth muscle and myometrial cell contractility, smooth muscle phenotype and vascular remodeling, and ocular neovascularization in the neonate (11–16). To discern the multifaceted SP-A-dependent and independent functions of SP-R210, we studied the basal phenotype of transgenic mice with conditional deletion of the SP-R210_L_ isoform. Our findings support the model that SP-R210_L_ modulates bi-directional communication between AMs and epithelial and vascular cells in the lung.

## RESULTS AND DISCUSSION

Selective deletion of SP-R210_L_ was achieved by Cre-mediated inversion recombination of a knock-in exon 1 allele with flanking Lox66/Lox77 recombination sites (Figure 1A) (17, 18). Breeding of *SP-R210_L_^fl/+^* with *Itgax(CD11c)^Cre^* mice (19) disrupted SP-R210_L_ but not SP-R210_S_ expression in AMs *SP-R210_L_^Invfl/+^* (Figure 1B-E). Exon 1 encodes the KE and PDZ domains of MYO18Aα. *Itgax^Cre^* targets mainly CD11c^hi^ AMs and dendritic cells (19). Immunofluorescence microscopy staining of lung tissue sections from *SP-R210_L_^fl/+^* littermate control mice(Figure 1F) and SP-R210_L_-deficient lungs (Figure 1H) localized SP-A to alveolar type II epithelial cells (yellow arrows), alveolar lumen, and bronchial club cells (white arrows) as previously established (20). SP-R210, however, localized to thin alveolar septa and the apical surface of dome shaped club cells in both control and SP-R210_L_-deficient lungs (Figure 1F-G, white arrows). SP-R210_L_ is the main isoform found in lung tissue and in stoichiometric expression with SP-R210_S_ in AMs (Figure 1C)(9). The overall fluorescence intensity for SP-R210 decreased by 20% in SP-R210_L_-deficient lungs compared to littermates, whereas SP-A staining was similar (Figure 1H). The number of AMs in aged SP-R210_L_-deficient mice decreased compared to littermates (Figure 1J), indicating AM attrition. The lung histology of SP-R210_L_-deficient mice did not show discernible abnormalities other than lymphocytic aggregates compared to littermates (Figure 1J,M). The SP-R210_L_-deficient AMs, however, stained positive for hemosiderin (Figure 1N-O) compared to no staining in controls (Figure 1K-L). Hemosiderin was confirmed in isolated SP-R210_L_-deficient AMs (Figure 1P-R), with highest levels in homozygous SP-R210_L_-deficient AMs (Figure 1S). BAL extracellular iron levels were not different (Figure 1T).

**Figure 1.**
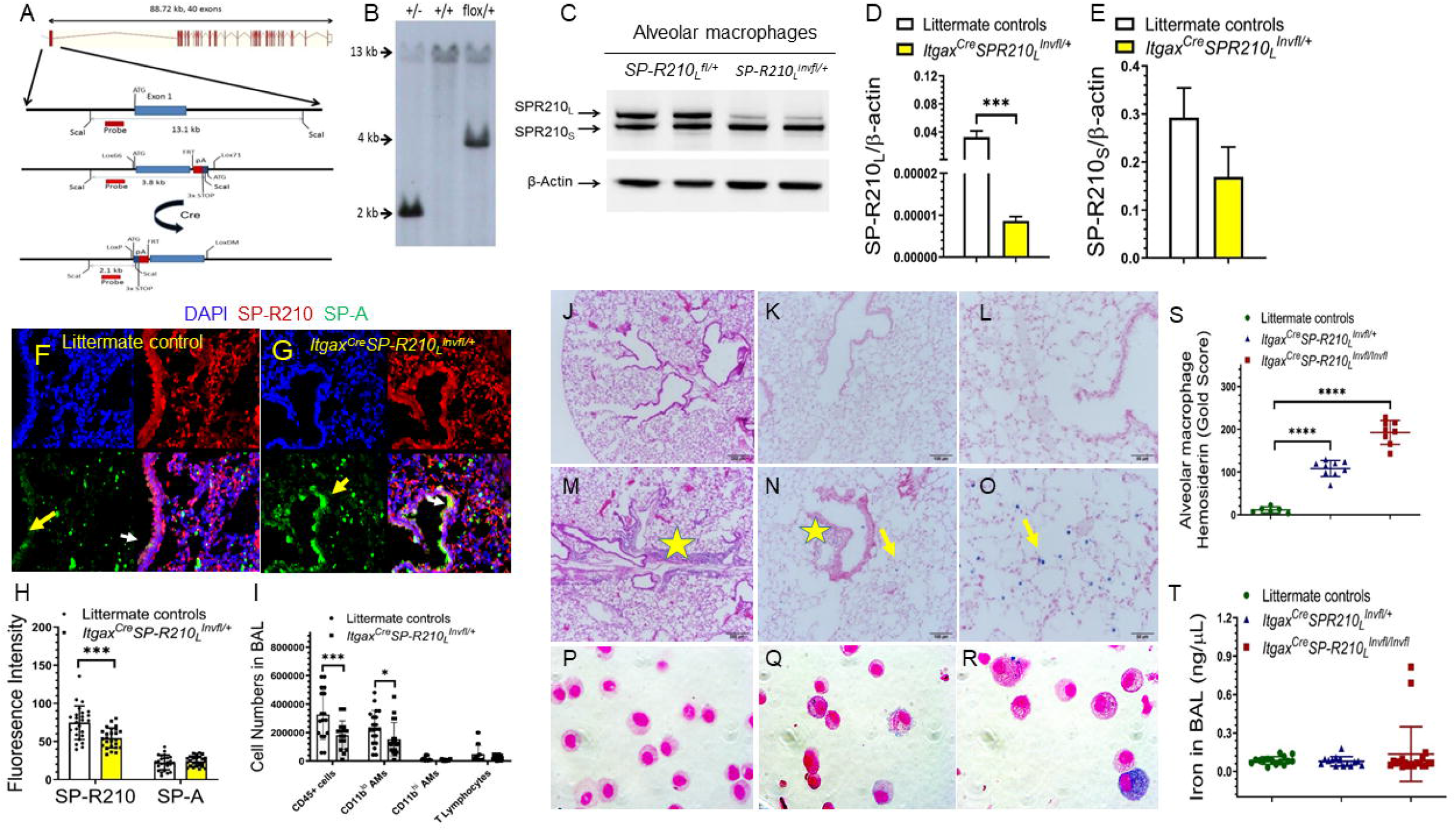
Disruption of SP-R210_L_ results in hemosiderin deposition in AMs and decline in AM number. A-B) Schematic depiction and targeting of the floxed *Myo18Aα* exon 1 by Cre-mediated inversion recombination. The targeted *SPR210_L_^Invfl/+^* allele was identified by Southern blot hybridization (B) and selective deletion of SP-R210_L_ protein was demonstrated by Western blotting (C) and densitometry analysis (D-E). D-E n=4 mice per group. ***p<0.001. F-G) Immunofluorescence microscopy and imaging of lung tissue sections showed similar distribution of SP-A and SP-R210 in control (F) and SP-R210_L_-deficient lungs (G) and partial reduction in SP-R210 fluorescence intensity (H) of SP-R210 in SP-R210_L_-deficient lungs compared to littermates. I) Flow cytometry analysis showed reduction CD45+ cells and CD11b^lo^ AMs cell number in BAL from aged 9-12-month old SP-R210_L_-deficient mice. J-O) H&E staining showed similar airway and alveolar histology of littermate (J) and SP-R210_L_-deficient lungs (M) and bronchocentric lymphocytic aggregates in the latter (yellow star in M). Perl’s Prussian blue staining revealed hemosiderin iron in SP-R210_L_-deficient AMs (N-O) but not littermate AMs (K-L). Perl’s iron staining of isolated AMs (P-R) confirmed lack of hemosiderin staining in littermate AMs (P) compared to hemosiderin AMs from heterozygous (Q) and homozygous (R) SP-R210_L_-deficient mice. S) Golde scoring showed proportional increase in hemosiderin ladden AMs in heterozygous and homozygous SP-R210_L_-deficient mice. For Golde scoring, n=8-12 mice per group at 8-11-months of age. ****p<0.0001. K) Similar concentration of free iron in BAL control and SP-R210_L_-deficient AMs.

Macrophage hemosiderin formation is part of normal iron flux and not readily detectable (21, 22). Under pathological conditions, AM hemosiderin develops from incomplete lysosomal degradation of ferritin, heme iron proteins, and partial hemoglobin degradation following erythrophagocytosis (23). SP-R210_L_-deficient mice did not develop abnormalities in systemic erythrocyte metabolism as indicated by similar hemosiderin spleen deposition (Figure 2A), hemoglobin degradation (Figure 2B), and kidney function (Figure 2C-D). Blood urea nitrogen levels (BUN) were similar at 2-months of age, although lower than controls in 6-months-old SP-R210_L_-deficient mice (Figure 2D). The aged SP-R210_L_-deficient mice did not develop tachypnea or dyspnea (Figure 2E). Basal capillary permeability was not affected (Figure 2F) as indicated by similar extravasation of Evans blue in the lung, although the circulating Evans blue concentration in SP-R210_L_-deficient mouse blood was 10% lower than littermates (Figure 2G), suggesting increased capillary volume. The lungs of 6-month-old SP-R210_L_-deficient mice, however, retained more fluid during heart perfusion compared to littermates (Figure 2H), indicating increased water permeability at increased pressure.

**Figure 2.**
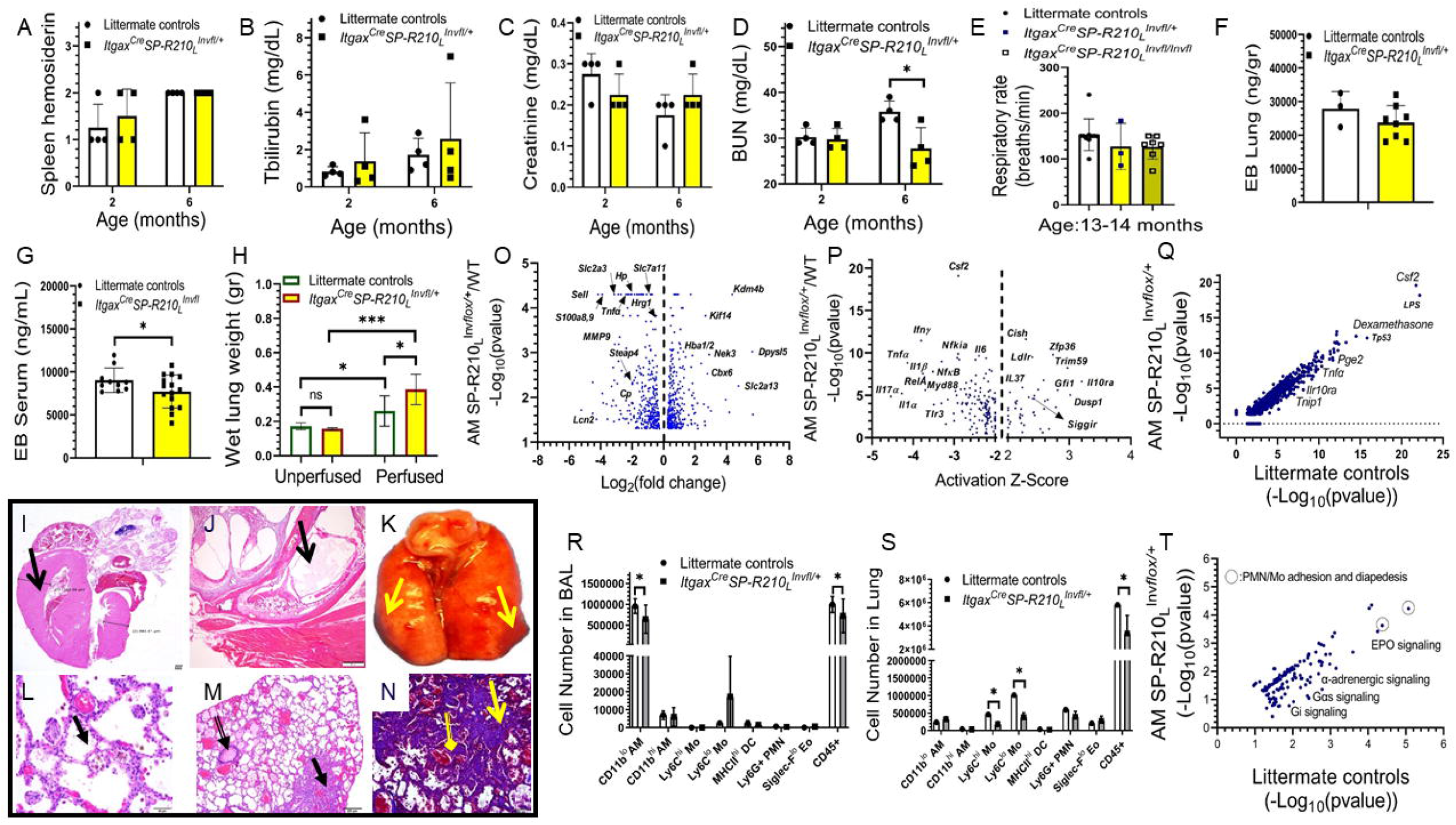
Disruption of SP-R210_L_ alters lung hydrostatic solute distribution, AM transcriptome phenotype, basal extravasation of monocyte subsets in the lung, and is associated with rare cardiopulmonary interstitial disease. . Histological scoring (A) and blood measurements (B-D) showed similar hemosiderin deposition in spleens of 2- and 6-momth old control and heterozygous SP-R210_L_-deficient mice as well as similar levels of T-bilirubin (B) and creatinine (C) in blood, whereas blood urea nitrogen (D) did not increase with age in SP-R210_L_-deficient mice. N=4 mice per group *p<0.05. Respiratory rates were also not different (E). F-H) Evans blue (EB) dye permeability experiments showed similar lung microvascular permeability of EB in 6-month old SP-R210_L_-deficient mice compared to littermates (F), whereas the concentration of circulating EB was lower (n=4, *p<0.05). Perfused SP-R210_L_-deficient lungs retained more fluid compared to littermates (n=6 per genotype in the unperfused group, and n=8 mice per genotype in the perfused group*p<0.05, ***p<0.001). I-N) Development of pulmonary hypertension (I), otitis media (J), lung hemorrhage (K), and intra-alveolar and interstitial fibrosis vasculopathy in a SP-R210_L_-deficient mouse (L-N) as detailed in “Results and Discussion”. (O-Q) Differential transcriptome profile (O) and upstream pathway analysis of SP-R210_L_-deficient AMs vs littermates (P-Q) indicate decreased threshold of basal AM activation. (R-S) Flow cytometry analysis showed decreased number of CD45+ cells and CD11b^lo^ AMs in BAL (R) and decreased number of both CD45+ immune cells and monocyte subsets (S) in lungs of 4-month old SP-R210_L_-deficient mice compared to littermates. (R-S, n=6 littermates and n=12 for *Itgax^Cre^SPR210_L_^Invfl/+^*, p<0.05, ***p<0.001)

Adding to this underlying pathophysiology, a 6-week-old female mouse presenting with lethargy and anorexia suffered from severe cardiopulmonary disease and otitis media (Figure 2I-N). Histological evaluation showed right ventricular and asymmetric septal hypertrophy (Figure 2I), unilateral otitis media with intraluminal serous exudate and neutrophilic inflammation affecting mainly the middle ear, and fibroplasia thickening of the ear canal lining (Figure 2J). In this regard, disruption of TGIF an inhibitor of TGFβ activation, a known effector of SP-R210 (24), leads to chronic otitis media (25). This case coincided with the presence of *Klebsiella oxytoca* and *Pseudomonas aeruginosa* in the mouse colony, suggesting an infectious trigger. Gross evaluation showed parenchymal lung hemorrhage (Figure 2K). Histopathological evaluation showed multifocal to coalescing expansion of alveolar septa with abundant nascent and mature fibrosis, abundance of foamy and frequently hemosiderin-laden intra-alveolar macrophages and fibrous projections, eosinophilic debris, and acute hemorrhage (Figure 2L-M). Mason’s trichrome staining showed fibrotic thickening of alveolar septa (Figure 2N). Small and medium arteries were dilated, tortuous, and redundant, with smooth muscle hyperplasia in smaller arteries (Figure 2L-N).

Comparative transcriptome profile analysis between SP-R210_L_-deficient and littermate AMs (Figure 2O-P) showed moderate increase in hemoglobin mRNAs, *Hba1, Hba2, and Hbb*, supporting erythrophagocytosis as the source of hemosiderin (Figure 2O) (26). Relevant to iron metabolism, RNAsequencing showed moderate decreases in heme and iron transport protein mRNAs for *Hrg1 (Slc48a1)* (27), ceruloplasmin (*Cp*) (28), *Steap4* (29), haptoglobin (*Hp*) (30), and lipocalin 2 *(Lcn2)* (31), indicating reduced iron import and intrinsic iron flux in SP-R210_L_-deficient AMs, responses consistent with iron abundance. The iron and α-ketoglutarate-dependent histone lysine demethylase *Kdm4b* gene (32) was the most highly expressed mRNA in SP-R210_L_-deficient AMs (Figure 1O), indicating epigenetic adaptation. KDM4B modulates repressive and non-repressive chromatin balance (32). One noteworthy KDM4B effector is PPARγ, which is central to surfactant metabolism in AMs (33). Relevant triggers of KDM4B expression include hypoxia (32) and osmolarity (34), conditions encountered during gestation and transition from the hypoxia *in utero* to normoxia in postnatal life (35–39), and TGFβ(32). In the present context, AMs may sense changes in oncotic balance in the lung’s airway-capillary barrier.

Upstream pathway analysis of mRNA sequencing data revealed reciprocal inhibition of inflammatory *(Tnfa, Il1β, RelA, Myd88, Il-17a, Ifnγ)* and activation of antiinflammatory pathways *(Cish, Zfp36, Il-10ra, Gfi1, Dusp1, Il37, Siggir)* in SP-R210_L_-deficient AMs (Figure 2P). Suppression of the *Csf2* signaling pathway (Figure 2P-Q) predicts decreased responsiveness of SP-R210_L_-deficient AMs to GM-CSF. Among the upregulated pathways, the IL-37/SIGGIR pathway (Figure 2P) can antagonize GM-CSF signaling (40). Relevant to the present findings, IL-37 is a proangiogenic (41) cytokine produced primarily by monocytes and dendritic cells (42). Furthermore, GM-CSF antagonizes SP-A binding and alters SP-A binding behavior in a concentration-dependent manner (43, 44). Conversely, SP-A maintains stoichiometric balance of SP-R210_L_ and SP-R210_S_ on AMs (2), modulates GM-CSF secretion (45), and has increased potency at suppressing TNFα in SP-R210_L_-deficient macrophages (2). The reduction in *Csf2* was associated with decreased AM number in both 4-month-old (Figure 2R) and aged SP-R210_L_-deficient mice compared to littermates (Figure 1I above). The number of lung CD45+ cells and monocytes were also suppressed (Figure 2S). To this end, monocytic recruitment genes (e.g. *Sell*, *S100A8/9, Cxcr2, TNFα)* were decreased (Figure 2O). Furthermore, canonical pathway analysis showed reduction in adhesion and diapedesis pathways in SP-R210_L_-deficient AMs (Figure 2T). Conversely, reduction in α-adrenergic receptor signaling (Figure 2S), suggested decline in neuro-hormonal control of SP-R210_L_-deficient AMs. In this context, cardiovascular autonomic (Figure 3A-H), arterial oxygen (Figure 3I), and hematological measurements (Figure 3J-P) did not support development of cardiac dysfunction in aging SP-R210_L_-deficient mice.

**Figure 3.**
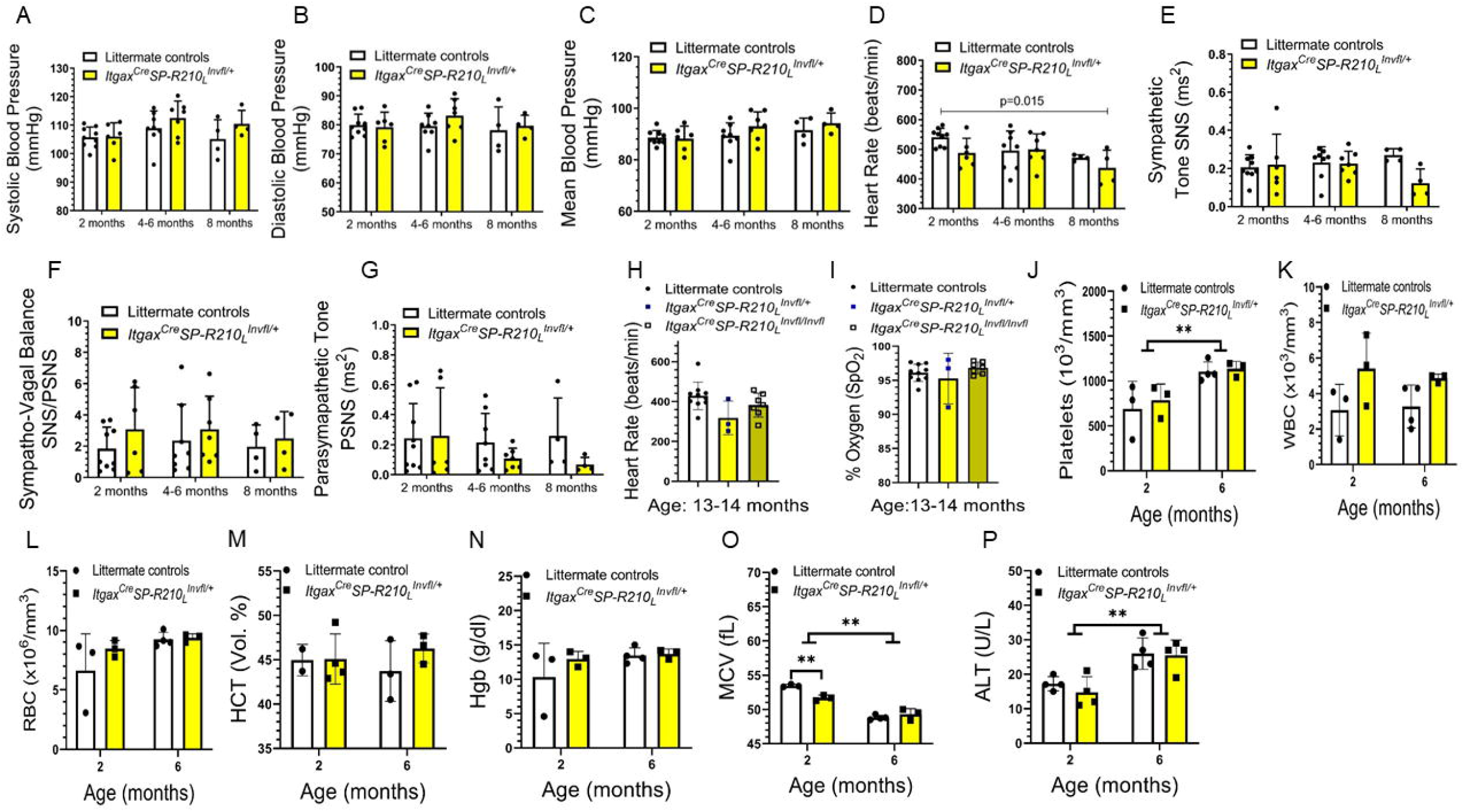
Disruption of SP-R210_L_ does not alter cardiac function, respiratory gas exchange, or hematologic parameters during aging. . Blood pressure (A-C), heart rate (D) and cardiac autonomic tone (E-G) were obtained under isoflurane anesthesia (n=6 mice/group at 2- and 6-months of age, and n=4/group at 8-months of age). Heart rate (H) and arterial oxygen saturation (I) in aged 13-14-month-old littermate (n=10), heterozygous (n=3), and homozygous (n=8) SP-R210_L_-deficient mice were measured using a MouseOx pulse oximeter under ketamine/xylazine anesthesia. Hematological parameters (J-O) and serum chemistry (P) were determined in blood collected via the cauda vena cava under ketamine/xylazine anesthesia. N=4 mice/group. All methods are described in “Supplemental Materials and Methods” and results detailed in “Results and Discussion”.

We next assessed if SP-R210_L_ deletion alters respiratory mechanics. Pressurevolume curves showed that the lungs of 6-month-old SP-R210_L_-deficient mice were less distensible (Figure 4A) and had increased expiratory curve slope (Figure 4B), indicating development of chronic lung disease. The age-associated increase in Inspiratory capacity (Figure 4C) and compliance (Figure 4D) in SP-R210_L_-deficient mice was slower than in littermates. Respiratory system (Figure 2E) and tissue elastance (Figure 2F) were not different. Central airway Newtonian resistance, however, increased over time in SP-R210_L_-deficient mice (Figure 4G), whereas tissue damping was not affected (Figure 4I). Respiratory system resistance remained stable in SP-R210_L_-deficient mice compared to decline in littermates (Figure 4I). All age-related changes in respiratory mechanics measured in control littermates reproduce previous studies in WT mice (46). Methacholine challenge studies showed enhanced airway responsiveness for respiratory system and tissue elastance (Figure 4J-K) and Newtonian resistance (Figure 4L) in 6-month-old SP-R210_L_-deficient mice compared to littermates. Tissue damping (Figure 4M) and respiratory system resistance (Figure 4N) where not different.

**Figure 4.**
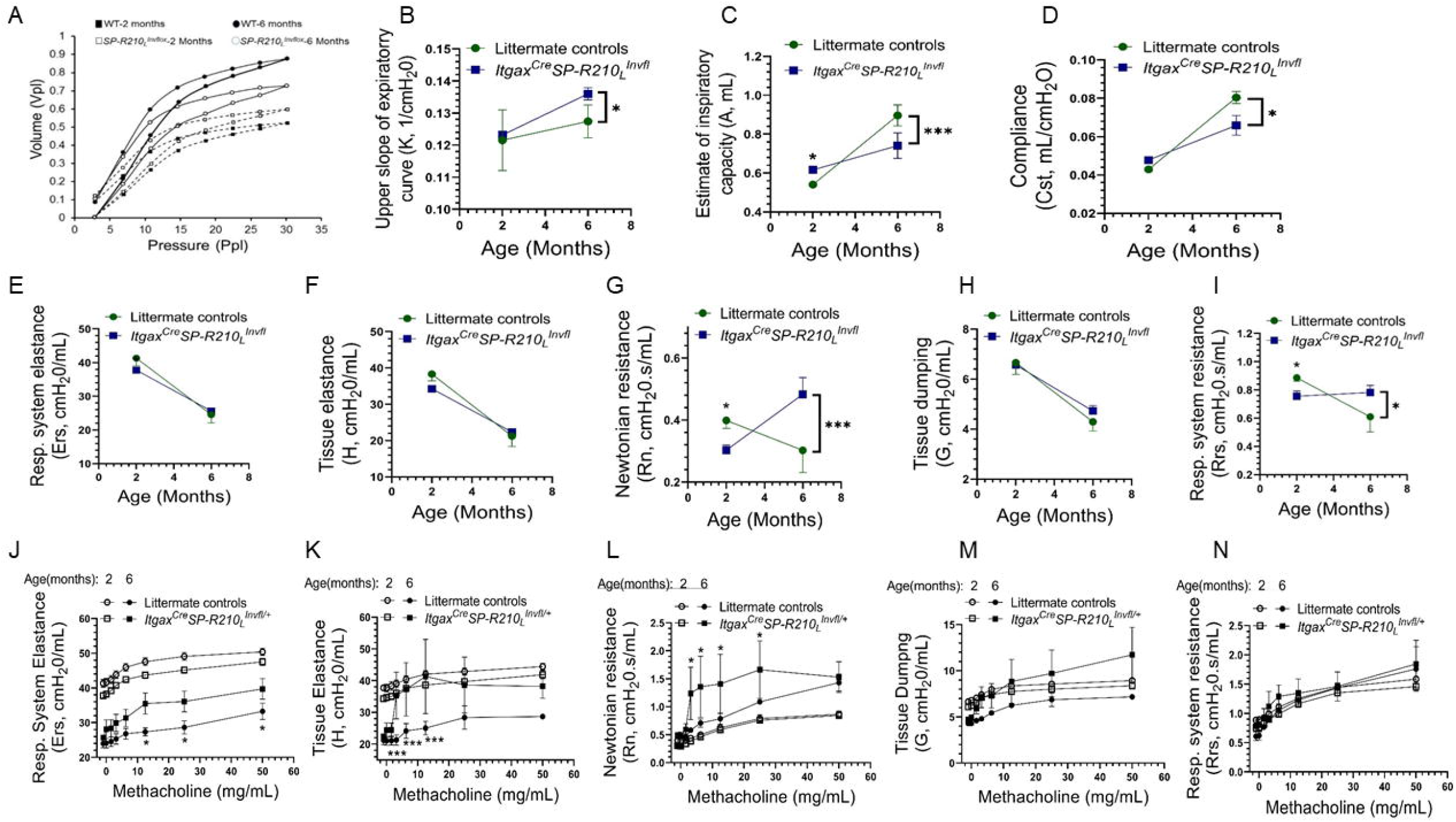
Disruption of SP-R210_L_ results in age-dependent alterations in basal respiratory mechanics and airway hyper responsiveness. Respiratory mechanics parameters were obtained using the forced oscillation technique as detailed in “Supplemental Materials and Methods” and “Results and Discussion” sections. N=4 mice per genotype per time point. *p<0.05, ***p<0.001.

Taken together, our findings indicate that SP-R210 isoforms modulate intercellular communication in the lung. Transcriptome data support the model that SP-R210 isoforms modulate rheostatic functions of SP-A and GM-CSF in the lung through epigenetic adaptation to local conditions. Selective disruption of the SP-R210_L_ isoform resulted in subclinical disease presenting as hemosiderin deposition, attrition of AMs, and changes in airway-capillary mechanics and basal fluid homeostasis in the lung. Rheological adaptation as indicated by differences in blood urea, serum Evans blue dye concentration, and hydrostatic fluid retention in older SP-R210_L_-deficient mice suggests compensatory oncotic fluid distribution to maintain respiratory function. Our study raises new questions about the mechanisms by which SP-R210 isoforms regulate immune and surfactant homeostasis, host defense, and respiratory physiology during development and aging.

## METHODS

The conditional *Myo18α* knock-in mice, designated as *SP-R210_L_^fl/+^*, were generated by Ozgene Pty Ltd. (Bentley WA, Australia) on a C57BL/6 background by homologous recombination. A knock-in exon 1 allele flanked by Lox66/Lox77 inversion recombination sites was made using standard molecular cloning techniques and transfected by electroporation in C57BL/6 embryonic stem cells (ES) cells. Homologous recombinant ES cells identified by Southern blotting were microinjected into C57BL/6 blastocysts. Founder transgenic were bred to establish the *SP-R210_L_^flox/+^* mouse colony and then with *Itgax^Cre^* mice for conditional deletion of SP-R210_L_ as detailed in “Supplemental Material and Methods”. All procedures were approved by the Institutional Animal Care and Use Committee of Pennsylvania State University College of Medicine and adhered to the 8^th^ edition of the PHS NIH GUIDE for the care and use of laboratory animals.

## AUTHOR CONTRIBUTIONS

EY, TMU, RA, KF, ZG, SH, SD, KH, and SB conducted experiments, acquired and graphed data, and edited the manuscript; EY processed and analyzed genomic data; HA and TKC conducted pathological diagnosis and blind scoring of histopathology; ACA and ESH designed cardiovascular and flow cytometry experiments, analyzed data and edited the manuscript; ZCC led and designed the study, analyzed data, and wrote the manuscript

## ACKNOWLEDGEMENTS

This work was funded by PHS grants HL128746, Pennsylvania Department of Health, The Children’s Miracle Network, and the Department of Pediatrics Pennsylvania State University College of Medicine. We would like to thank Nate Schaffer and Joseph Bednarzyk from the Penn State College of Medicine Flow Cytometry Core Facility as well as the Institute of Personalized Medicine for assistance with genomic processes.

