## Supplemental Figures and Tables for "Disruption of the surfactant protein A receptor SP-R210_L_ (CD245α/MYO18Aα) alters respiratory function and iron sequestration in alveolar macrophages of aged mice"

### SUPPLEMEMENTAL MATERIALS AND METHODS

*Generation of conditional SP-R210$_L$-deficient mice*

The conditional *Myo18α* knock-in mice, henceforth designated as *SP-R210$_L^{fl/+}$*, were generated by Ozgene Pty Ltd. (Bentley WA, Australia) on a C57BL/6 background by standard homologous recombination. A knock-in allele flanked by Lox66/Lox77 inversion recombination sites was made using standard molecular cloning techniques and transfected by electroporation in C57BL/6 embryonic stem cells (ES) cells. Homologous recombinant ES cells were identified by Southern blot analysis and microinjected into C57BL/6 blastocysts. Offspring were backcrossed to C57BL/6 mice and germline transmission was confirmed by Southern blot analysis of tail genomic DNA (1-3) (Figure 1A-B). The *SP-R210$_L^{fl/+}$* mice were crossed to hemizygous Tg(Itgax-Cre)1-1Reiz (*Itgax$^{Cre}$*) mice (4) to obtain *Itgax$^{Cre}$SP-R210$_L^{Invfl/+}$* mice carrying the heterozygous inverted allele and wild type (WT) and *SP-R210$_L^{fl/+}$* littermate controls. *Itgax$^{Cre}$SP-R210$_L^{Invfl/+}$* mice with paternal transmission of *Itgax$^{Cre}$* were then mated to WT or *SP-R210$_L^{fl/+}$* mice to obtain heterozygous *SP-R210$_L^{Invfl/+}$* mice and these were mated to each other to obtain homozygous *SP-R210$_L^{Invfl/Invfl}$* mice and littermate controls. The SP-R210$_L$ deletion did not alter embryonic survival, fecundity, or lifespan. *Itgax(CD11c)$^{Cre}$* targets genes mainly in CD11c$^{hi}$ macrophages and dendritic cells (4, 5) but can also target genes in monocyte, neutrophil, and lymphoid cell subsets (5, 6). Alveolar type II epithelial cells and club cells can also be targeted as they express moderate to abundant *Itgax* mRNA (7), whereas low level expression of cre in germ cells exerts deletion of floxed genes that are highly sensitive to cre-mediated recombination (6). Inversion recombination and deletion of SP-R210$_L$ was confirmed by Southern blot hybridization of tail DNA biopsies and western blot analyses of alveolar macrophages (AMs), respectively (Figure 1B-C). The present study utilized 2-month and ≥ 6 months old *Itgax*$^{Cre}$*SP-R210*$_L^{Invfl/+}$, *SP-R210*$_L^{Invfl/+}$, and *SP-R210*$_L^{Invfl/Invfl}$ male and female mice designated collectively as *Itgax*$^{Cre}$*SP-R210*$_L^{Invfl/+}$ or *Itgax*$^{Cre}$*SP-R210*$_L^{Invfl/Invfl}$ in most figures. All procedures were approved by the Institutional Animal Care and Use Committee of Pennsylvania State University College of Medicine and adhered to the 8$^{th}$ edition of the PHS NIH GUIDE for the care and use of laboratory animals.

*Mouse genomic DNA isolation*

Tail snips from 3-week-old mouse pups were digested using 300 µL of 100 mM Tris, 5 mM EDTA, pH 8.0, 200 mM NaCl, 0.2% sodium dodecyl sulfate (SDS) and 0.4 mg/mL Proteinase-K overnight at 55$^o$C. The DNA was precipitated with 1 mL of 100% ethanol and centrifuged at 16,000 x g for 30 minutes at 4$^o$C. DNA precipitates were washed with 1 mL of 70% ethanol and centrifuged at 16,000 x g for 20 minutes at 4$^o$C. The DNA pellets were then dissolved in 60 µL of 10 mM Tris-HCl, 0.2 mM $Na_2EDTA$, pH 7.5 and the excess ethanol was evaporated at 55$^o$C for approximately 2 h. Isolated DNA was stored at -20$^o$C.

*Southern blot hybridization genotyping*

Digoxigenin (DIG)-labeled DNA probes were synthesized using the Roche DIG synthesis kit (SIGMA, Cat no 11636090910) in a 50 µL reaction mix consisting of 100 pg plasmid DNA template, 5 µL 10X buffer, 5 µL DIG mix, 10 µM each of (5’-TTCAGAGGGGAGGCATTCATGGAG-3’) and (5’-GATGTGTCTTATCGGGAGCTTCAC-3’) PCR probes, and 0.75 µL of Expand High Fidelity Polymerase. The PCR reaction was denatured for 2.0 min at 95$^{o}$C followed by 40 cycles at 95$^{o}$C 30 min, 60$^{o}$C 30 sec, 72$^{o}$C 40 sec, and final elongation at 72$^{o}$C for 7 min. The 559 bp product was gel extracted and stored at -20$^{o}$C till use. Genomic DNA was digested overnight at 37$^{o}$C in a 20 μl reaction mix containing 8 υμg of DNA, 0.5 units of ScaI-HF (New England Biolabs), 2 μL of 10X Cutsmart buffer. Digested DNA and Digoxigenin-labeled DNA molecular weight standards (Roche Diagnostics, Cat no. 11218590910) were loaded onto 0.7% agarose gels prepared in 0.5X TBE buffer and separated for 2.5 h at 150V. Gels were immersed in denaturation solution containing 0.5 M NaOH and 1.5 M NaCL two times for 15 min with shaking. Denatured gels were then rinsed in dd$H_2O$, submerged in neutralization solution containing 0.5 M Tris-HCl, pH 7.5 and 1.5 M NaCl two times for 10 min and the DNA was then transferred onto charged nitrocellulose in 20X SSC by capillary action. Transferred DNA was fixed by UV crosslinking for 1-3 min. The nitrocellulose membranes were then rinsed in dd$H_2O$ and incubated in preheated DIG Easy hyb buffer for 15-30 min at 42$^{o}$Cand then annealed with 5-25 μg/mL of DIG-labeled DNA probe in DIG Easy Hyb buffer (3.5ml/100cm$^2$ membrane) with gentle agitation overnight at 42$^{o}$C. Hybridized membranes were then washed three times for 10 min in 1 X SSC, 0.1% SDS for 10 min at room temperature with agitation and then two times for 15 min at in 0.2 X SSC, 0.1% SDS for 15 min at 68$^{o}$C under constant agitation. Hybridized membranes were washed in DIG Wash solution for 1-5 min and blocked in DIG Block Buffer (SIGMA, Cat no 11585762001) for 1 hr. Hybridized DIG-labeled DNA was visualized on X-ray film using the Ready-to-use CDP-Star (Cat no 12041677001).

*PCR genotyping*

Mice carrying WT (*SP-R210*$_L^{+}$), floxed (*SP-R210*$_L^{fl}$) and inverted knockout (*SP-R210*$_L^{Invfl}$) alleles were identified by PCR amplification of genomic DNA using three sets of primers. The sequence of the forward and reverse primers were 5' GTC TGA GGT TCC CTT CAA TGCC-3' and 5'-TCC TAT ACT CAC AGC TTC TCA G for WT, 5'-ACA ACA GAT GGC TGG CAA CTA GA-3' and 5'- TCC TAT ACT CAC AGC TTC TCA G-3' for the floxed allele, and 5-ACA ACA GAT GGC TGG CAA CTA GA-3' and 5'-AAG GTC TTT GAG GCA GGG GCT AAG-3' for the inverted knockout allele. The *Itgax*$^{Cre}$ transgene was identified using 5'-ACT TGG CAG CTG TCT CCA AG-3',5'-GCG AAC ATC TTC AGG TTC TG-3', as primers and internal control primers 5'-CAA ATG TTG CTT GTC TGG TG-3' and 5'-GTC AGT CGA GTG CAC AGT TT-3' resulting in 313 and 200 bp PCR amplicons for the transgene and internal controls respectively. PCR reactions were carried out in a volume of 25 μL containing 100 ng of genomic DNA template, 12.5 μL of 2X Econo Taq® PLUS mix (Lucigen), and 10 μM of each primer. Three PCR reaction conditions were initial denaturation at 95°C for 5 min, followed by 35 cycles of denaturation at 95°C for 1 min, annealing at 58 °C for 1min, extension at 72°C for 1 min and a final elongation step at 72°C for 5 min. The reaction conditions for *Itgax*$^{Cre}$ genotyping were initial denaturation at 94°C for 3 min, followed by 35 cycles of denaturation at 94 °C for 30 sec, annealing at 64 °C for 1 min, extension at 72°C for 45 sec and a final elongation step at 72°C for 2 min. The PCR products were mixed with 2 μL of 6X loading dye (0.25% Bromophenol Blue, 0.25%Xylene Cyanol FF, 30% Glycerol), loaded onto 1% ethidium bromide stained agarose gels, and electrophoresed at 150 V for 5 mins. Gel images were recorded using a Bio-Rad ChemiDoc™ MP imaging system.

*Hemosiderin and iron measurements*

Alveolar macrophages were collected by bronchoalveolar lavage and deposited on glass slides by centrifugation at a density of 25,000 cells perslide using a Shandon cytospin centrifuge (Cytospin; Shandon Southern Instruments, Runcorn, UK) at 1000 rpm for 10 min. Perl's Prussian blue was used to stain hemosiderin. Air dried slides were hydrated in water and then placed in a stock 10% potassium ferrocyanide solution for 5 min, followed by an acid ferrocyanide solution consisting of 1% hydrochloric acid and7% potassium ferrocyanide for 20 min. Slides were then rinsed well in ddH2O, and counterstained with nuclear fast red stain consisting of 0.1% (w/v) in 5% aluminum sulfate containing thymol as a preservative. Stained slides were washed in lukewarm tap water for 10 min, mounted in Permount, and visualized under a microscope. Hemosiderin deposition was scored blindly on a scale of 0 to 4 (0 – no blue color; 1 – faint blue color stain in cytoplasm; 2 –dense blue color in minor portion of the cytoplasm or medium color intensity through cell; 3 – deep blue staining in most of cytoplasm; 4 – dark blue throughout the cytoplasm). The Golde hemosiderin score was obtained by dividing the total score from 200 cells by 2. After scale assignment, an average score was calculated for, zero being the minimum and 400 the maximum score (8, 9). The concentration of extracellular iron was measured using a commercial colorimetric assay (SIGMA, Cat no MAK025) in BAL supernatant after centrifugation at 16,000 x g for 10 mins.

*Respiratory mechanics*

Parameters of lung function were measured using the forced oscillation technique and a computer-controlled flexiVent FX ventilator (SCIREQ, Montreal, Canada) as previously described (10). Briefly, mice were anesthetized with a mixture of ketamine/xylazine and a 19G cannula was secured in the trachea. Mice were then connected to the flexiVent via the cannula and ventilation started in oxygen containing 1% isoflurane at a respiratory rate of 150 breadths per minute. The non-polarizing paralytic vecuronium bromide was administered at this time to block respiratory movements. After 5 min of ventilation, manual deep Inflation and manual pressure-volume loop (PVs-P) scans were performed to collect baseline parameters before proceeding with Methacholine inhalation dose response scans. methacholine (acetyl-β-methacholine chloride, Sigma-Aldrich, St Louis, MO) was freshly prepared in DPBS at doses of 0, 1.56, 3.13, 6.25, 12.5, 25 and 50 mg/mL and administered using the flexiVent Aeroneb fine particle nebulizer. The script used for the inhaled dose response included two Deep Inflation scans followed by twelve repeats of alternating SnapShot (sinusoidal – single frequency forced oscillation waveform) and Primewave (broadband – multi-frequency forced oscillation waveform) scans at baseline and for each methacholine dose. Data were then analyzed using flexiWare software (SCIREQ) and exported to Excel for further analyses.

*Respiratory rate and oxygenation*

Breath rate (brpm), breath distention (μm) and arterial oxygen saturation ($S_pO_2$) were measured using a MouseOx Plus pulse oximeter (STARR Life Sciences Corp., Oakmont, PA) and mouse thigh clip sensor. Mice were anesthetized using a mixture of ketamine/xylazine and placed on a heating pad to maintain body temperature. The right thigh of each mouse was shaved and the thigh clip sensor attached. Parameters were monitored using the MouseOx Plus software and data collection was started after all parameters had stabilized. Collected data were saved as text files for further analysis using Excel, excluding any data points flagged with error codes by the MouseOx Plus software.

*Hemodynamic measurements*

Blood pressure and heart rate were measured in isoflurane-anesthetized SP-R210$_L$-deficient mice and littermate controls at 2 months (n=6-9/group) and at 4 to 6 months (n=7-8/group) via an indwelling carotid artery catheter connected to a strain gauge transducer and blood pressure analyzer (MP36R, Biopac Systems, Inc., Goleta, CA, USA). Continuous recordings of blood pressure and heart rate were obtained for at least 10 min, reporting the average values across this time period. The blood pressure and heart rate recordings were analyzed using frequency domain spectral analysis methods to obtain indices of cardiac sympathetic and parasympathetic tone (Acqknowledge 4 software, Biopac Systems, Inc.) as previously reported (11).

*Microvascular permeability*

Microvascular permeability was measured using an Evans blue dye Assay as previously described (12). Briefly, mice were intraperitoneally injected with 4mL/kg body weight with a sterile-filtered solution of 1% Evans blue dye (w/v) in PBS. After 24 hours of circulation, mice were anesthetized using a mixture of Ketamine/Xylazine and samples collected.

Blood was collected from the caudal vena cava and placed into BD Vacutainer blood collection tubes (Becton Dickinson, Franklin Lakes, NJ) containing EDTA for plasma. Mice were then perfused transcardially with 50mL of PBS followed by 20mL of 10% neutral buffered formalin and the lungs collected and weighed. Lungs were lyophilized and extracted with formamide (2mL/g of tissue) at 65°C for 24 hours. If necessary, plasma and lung extracts were diluted with PBS and the concentration of Evans blue dye determined spectrophotometrically at 620nm against an Evans blue dye standard curve. Both plasma and tissue extract samples were corrected for heme by taking a second absorbance reading at 740nm: $A_{620(corrected)} = A_{620} - (1.426 \times A_{740} + 0.030)$.

*Hematology and blood chemistry*

Hematological and blood chemistry analyses were performed by the Comparative Medicine Diagnostic Laboratory at the Pennsylvania State University College of Medicine. Mice were anesthetized using a mixture of ketamine/xylazine and blood collected from the caudal vena cava. Blood was placed into bBD Vacutainer blood collection tubes containing EDTA for hematological analyses including complete blood count with differential (CBC with differential), and into non-EDTA containing tubes and allowed to clot for serum chemistries.

*Bacterial infection*

Mice were infected at 2 months of age via the intranasal route in 40 μL PBS containing 100,000 colony forming units of *Streptococcus pneumoniae* strain A66.1 provided by

David Briles (University of Alabama at Birmingham, Birmingham, AL) (13, 14) as previously described (15).

*Preparation and administration of IL-4c*

The IL-4c complex was prepared by incubation of 5 υg of murine recombinant IL-4 (Peprotech, 214-14) with 25 μg anti-murine IL-4 (BioXCell clone: 11B11) for 10 min on ice per dose followed by dilution to 100 μL in PBS per intraperitoneal injection as previously described (16). Each mouse was given IL-4c at -4 and -2 days and 1 mg of BrDU (Sigma) 3 h prior to harvest of alveolar macrophages by bronchoalveolar lavage and lung cell suspensions (16, 17). Cell suspensions were processed by flow cytometry and alveolar macrophages gated as CD11b-SiglecF+ and CD11b+Siglec-F+ cells in BAL and post-lavage lung cell suspensions as described previously (18) respectively. Proliferating and RELMα+ cells were identified by intracellular co-staining with Ki67 (BD Pharmingen Cat no 556026), BrDU antibodies (Biolegend, Cat no 339812), and RELMα (Peprotech, Cat no. 500-P214) antibodies.

*Bronchoalveolar lavage and lung cell collection*

Alveolar macrophages were isolated by bronchoalveolar lavage (BAL) as previously described (18). Anesthetized mice were intra-tracheally cannulated with Intramedic™ (BD Bioscience, Franklin Lakes, NJ) polyethylene tubing (ID: 0.58 mm, OD: 0.965 mm) and BAL cells collected in 2.5 mL DPBS total with 1 mM EDTA. Lungs were perfused via a single cardiac perfusion of 10 mL DPBS and then washed with one lavage of 1 mL HBSS with $Ca^{+2}$ and $Mg^{+2}$. Lungs were then infused with 1 mL of digest solution (50 μL Liberase DL (Roche Cat no 5466202001), 1X DNase I (Roche Cat no 10459001) per 6 mL of HBSS w/ $Ca^{+2}$ and $Mg^{+2}$). The intra-tracheal tube was slowly removed while tying the trachea shut, and then the lung was excised from the thoracic cavity. Lung lobes were separated from the trachea and bronchi and then minced. Minced tissue was incubated with 5 mL of digest solution for 30 min at 37°C, and then homogenized by repeatedly passing the tissue suspension through an 18G needle. The cell suspensions were incubated for an additional 15 more min at 37°C and then 8 mL of stop solution (PBS with 2% FBS, 5mM EDTA, pH8.0) was added and the entire suspension was passed through 40 μm filter; an additional 20 mL of stop solution was used to wash the filter. BAL and Lung homogenate samples were centrifuged at 1200 RPM for 7 min at 4°C and resuspended in 10 mL of cold wash buffer. Cells were spun down one more time and resuspended in 1 mL 1X erythrocyte lysis buffer (eBioscience Cat no 00-4300-54) for 7 min at room temperature. Samples were centrifuged and resuspended in 1 mL wash buffer.

*Pathology and Immunohistochemistry*

Lungs were inflated with 4% paraformaldehyde in PBS and then fixed overnight in 4% paraformaldehyde in PBS before transferring to 75% ethanol. Cross-sections were cut for the lobes of each lung and placed into a single cassette. Lung lobe cross-sections were embedded in paraffin with slides prepared by the Comparative Medicine Diagnostic Laboratory at the Pennsylvania State University College of Medicine. Slides were deparaffinized with xylene and rehydrated using decreasing percentages of ethanol (100,

95, 70, 50, 0). For pathological assessment, lung tissue sections were stained with hematoxylin and eosin or Perls' Prussian blue staining. Images were captured using Olympus BX51 microscope and DP71 digital camera using CellSens Standard 1.12 imaging software (Olympus America, Center Valley, PA). Sections of liver, spleen, and heart were reviewed for all possible microscopic lesions by a veterinary pathologist blinded to cohort. The spleen was scored for several lesions, including white or red pulp intra-histiocytic pigment (presumed hemosiderin), increased extramedullary hematopoiesis, and marginal zone or germinal center hyperplasia. Lesions were scored using a 0-4 scale where 0 equals no lesion; 1 refers to a lesion that is focal and minimally elevated above normal limits; 2 are notable, multifocal microscopic changes that are still mild; 3 are moderate changes that disrupt a significant portion of the tissue's organization; and 4 where there are marked, diffuse changes.

For immuno-fluorescent staining, background auto-fluorescence was reduced by washing slides three times ten min each with 1 mg/mL sodium borohydride in PBS. All steps for the immunostaining of slides were performed in Shandon Sequenza trays (Thermo Fisher Scientific, Waltham, MA). Briefly, slides were blocked at room temperature with 5% horse Serum, 1% BSA, 0.1% Triton-X100, 1:500 Purified Rat Anti-Mouse CD16/CD32 mouse Fc block (BD Biosciences clone 2.4G1, cat no: 553142) and primary antibody incubations were done overnight at 4°C in blocking solution. Slides were then washed with PBS and secondary antibody incubations done for one hour at room temperature in blocking solution. Slides were again washed with PBS, briefly rinsed with water and coverslipped with Prolong Diamond Antifade Mount with DAPI (Thermo Fisher Scientific). Images were captured with a Leica DM4000 B LED microscope using the

Leica Application Suite X (LAS X) 3.0.0.15697. Average fluorescence intensity was acquired using the Stack Profile function in LAS X. Average fluorescence for 5 images per lung section for each of the five lobes per genotype were obtained and analyzed. Primary antibodies used included: Rabbit anti-SP-A (19) and mouse anti-SP-R210 monoclonal antibody (IgG2b) (clone P1H3 prepared in house) (20), Secondary antibodies used included: AF647 goat anti-rabbit IgG (ThermoFisher, cat no: A-21244) and AF594 AffiniPure donkey anti-mouse (Jackson ImmunoResearch, cat no: 715-585-150).

*RNA sequencing analysis*

RNA was isolated from alveolar macrophages collected by bronchoalveolar lavage using Trizol as described previously (18). RNA concentration and purity were determined by NanoDrop and A260/A280. RNA libraries were synthesized via single stranded poly-A isolation and sequenced at a depth of 25M as detailed previously (17). Sequences were aligned to the mm10 reference genome using hisat2 and RPKM values were obtained using featurecounts and edgeR. RPKM values were placed into Ingenuity pathway analysis (IPA, www.qiagen.com/ingenuity) to obtain fold expression differences. List of differentially expressed genes with a p-value of less than 0.05 were exported and futher analyzed. Genes were filtered against the MGI database of immune-associated genes. Genes were mapped to KEGG pathways using fgsea.

*Statistics and graphical analysis of data*

All statistical analysis were performed using tools embedded in GraphPad Prism software version 9.0. Main effects of age and genotype, and their interaction, were assessed by two way ANOVA followed by Tukey's post-hoc test ($\alpha$=0.05). Group comparisons were performed using the multiple t-test and p values calculated using the Holm-Sidak method. Differences were considered significant at $p<0.05$. Survival analysis was performed using Kaplan –Meier and curves and comparisons were based on the log-rank test.
